# Multimodal 3D light-field and laser-speckle endoscopy

**DOI:** 10.64898/2026.06.30.735698

**Authors:** Corey Zheng, Shu Jia

## Abstract

Minimally invasive surgery is a powerful technique that enables operations deep within the body while minimizing patient trauma and recovery time. Optical endoscopes are key to providing intraoperative vision but still face challenges due to the loss of essential senses, including depth perception and tactile feedback for tissue evaluation. Thus, it is critical to develop endoscopic imaging technologies that can augment operators with critical information. In this work, we explore a prototype multimodal 3D imaging endoscope that integrates volumetric light-field imaging with laser-speckle contrast imaging to simultaneously capture 3D structure and blood-flow information in a clinically relevant form factor.

## 1. Introduction

Optical endoscopy is a cornerstone of modern minimally invasive surgery (MIS), enabling clinicians to visualize and operate on internal tissues through small incisions, minimizing patient trauma. Owing to these advantages, MIS has become standard practice across nearly all interventional specialties [1]. However, the transition from open surgery to MIS introduces significant perceptual challenges for the operator. Endoscopic visualization is typically monocular, providing only a two-dimensional projection of the operating field. Furthermore, without direct access to the surgical site, there is no tactile feedback, which is commonly used to assess tissue viability and perfusion [2, 3]. Together, the loss of stereoscopic depth cues and the inability to assess tissue viability can compromise the surgeon’s spatial awareness and knowledge of the patient’s condition, contributing to intraoperative uncertainty.

Recently, stereo endoscopes have been developed to partially restore depth perception by integrating two image sensors into the endoscope tip and presenting each eye with a separate view through polarized displays [4]. While this approach recreates binocular disparity [5] and has been shown to improve surgical performance in some tasks [6, 7], it does not capture all three-dimensional binocular cues, and operators have reported adverse effects, including fatigue and motion sickness, leading to mixed clinical reception [4, 7, 8]. Furthermore, stereo endoscopes do not directly produce quantitative volumetric data, which may necessitate additional preoperative imaging, such as computed tomography (CT), to obtain the 3D structure of the tissue [9]. Such quantitative volumetric information is increasingly essential for surgical path planning and the emerging field of autonomous robotic surgery [10, 11].

Comparatively, light-field (LF) 3D imaging systems have shown promise for surgical applications. An emerging technology for surgical applications [12–14], light-field imagers, compared with stereo-based systems, utilize three or more views of the scene, typically by placing a micro-lens array in front of the camera [15]. This results in several advantages over stereo imaging, including reduced calibration requirements and the ability to image disparity along both the vertical and horizontal axes [16], leading to improved 3D reconstruction accuracy [11] and enabling renderings of 3D volumes rather than relying on stereo perception. As a result, they have been successfully applied to autonomous surgical robotics [17] and endoscopy [18, 19]. Furthermore, light-field imaging can be fused with several common computational approaches for 3D reconstruction [20], including structure-from-motion (SfM) and simultaneous localization and mapping (SLAM), thereby enhancing their performance [21, 22]. However, light-field endoscopes still require further development, including optical and form factor improvements, to meet clinical needs [18, 19].

Beyond structural visualization, there is a critical and largely unmet clinical need for continuous, non-invasive monitoring of blood perfusion during surgery. Tissue viability assessment is essential in procedures such as bowel anastomosis, neurosurgery, and free flap reconstruction, where inadequate perfusion can lead to ischemia and tissue necrosis. Current clinical practice relies primarily on dye-based angiography using agents such as indocyanine green or fluorescein, which require intravenous injection, provide only temporary visualization, and cannot support continuous monitoring [23–25]. Laser speckle contrast imaging (LSCI) is a promising alternative that utilizes changes in a dynamic speckle pattern caused by the motion of the scattering medium [26]. Enabling the widefield computation of relative blood flow maps efficiently using straightforward instrumentation, LSCI has been demonstrated in open neurosurgical and dermatological settings [27] and has been integrated with monocular endoscopes [28, 29], but has seldom been investigated in combination with three-dimensional imaging [30].

The unification of light-field volumetric imaging and laser speckle contrast analysis in a single endoscopic device would provide surgeons with simultaneous access to tissue surface topography and perfusion. However, three-dimensional laser speckle imaging has yet to be addressed in a clinically viable form factor [31]. Here, we introduce a multimodal 3D imaging endoscope that integrates light-field volumetric reconstruction with laser speckle contrast perfusion mapping and investigate its potential as a tool for intraoperative care. This design presents a significantly improved optical design compared to our previous works[18, 19], and we demonstrate the additional capability of the system to extract relevant metrics of blood flow in a 3D structure, proving the compatibility of laser-speckle contrast imaging with light-field reconstruction techniques. We further evaluate the system’s 3D imaging performance across a range of targets, from inanimate objects to tissue phantoms and real biological tissues, and explore its performance under different operating conditions.

## 2. Results

### 2.1 System overview

The multimodal endoscope system is overviewed in Figure 1. The endoscope comprises seven gradient-index (GRIN) lenses arranged in a hexagonal array. Conceptually, this light-field lens array allows changes in the depth of any point within the field of view (FOV) to be encoded as lateral translation in the image across different perspective views (Figure 1A, dotted inset). A relay system re-images the intermediate images from the GRIN lens array onto the cameras, providing aberration correction and magnification. Using a dichroic mirror, a dual-imaging and illumination system captures both regular RGB and near-infrared (NIR) images (Figure 2A). The NIR imaging channel is used to capture speckle dynamics from a coherent 785nm source, which can be processed into blood flow-index maps [32]. Then, using the calibrated point-spread functions (PSFs) of the system (Figure 1A, solid inset), both the RGB light-field image and the computed light-field flow maps are volumetrically reconstructed using Richardson-Lucy Deconvolution (RLD), yielding 3D volumetric information for both structure and flow. Compared to previous works, our as-fabricated system presents a more clinically relevant form factor [18, 19], with a working length of over 100 mm with a 5.2 mm diameter, comparable to other existing endoscopic technologies [33].

**Figure 1.**
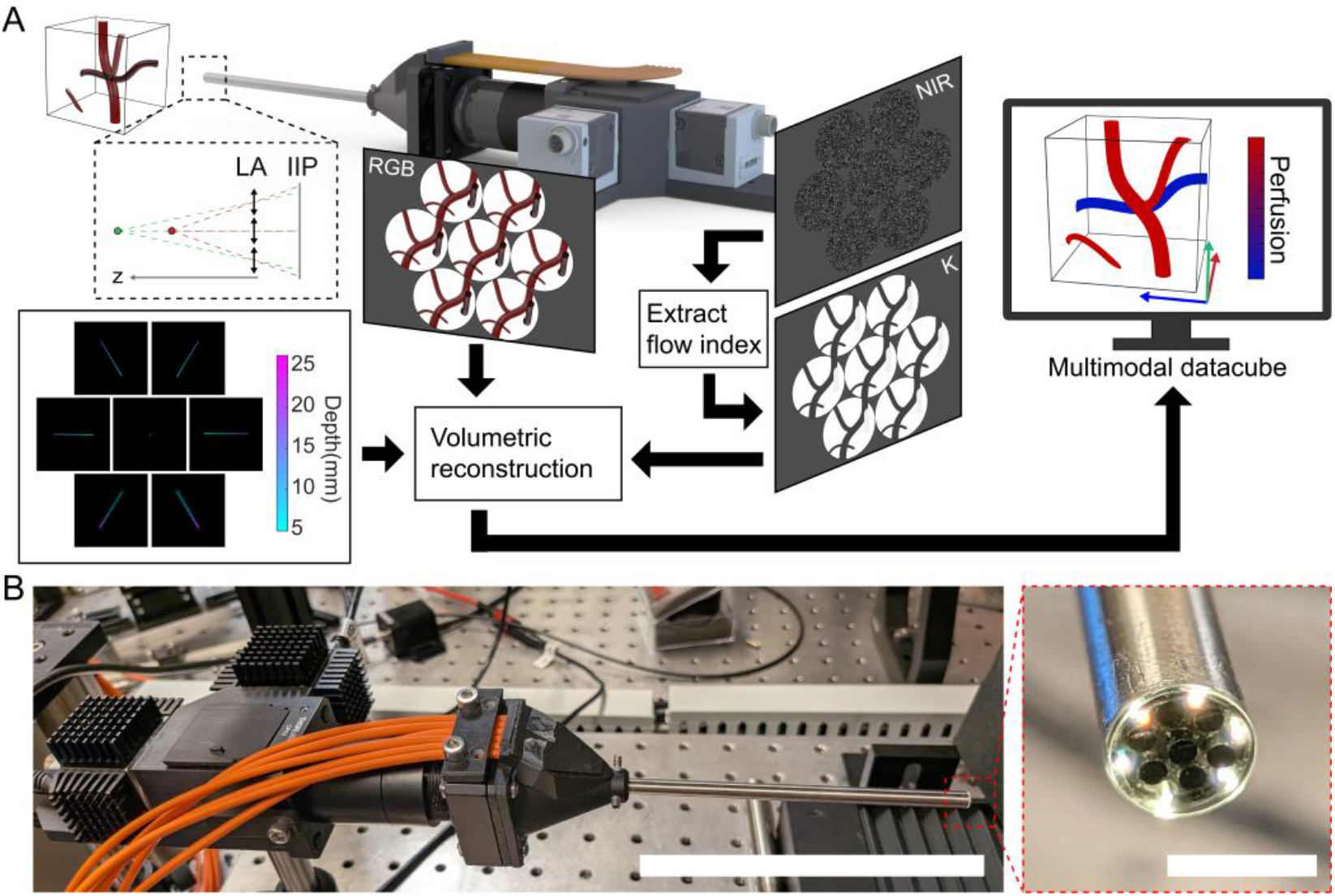
System overview of multimodal endoscope. (A) Operating principle, showing imaging, reconstruction, and final output flowchart. Dotted inset: schematic illustrating the ability of a lens array (LA) to encode depth information as a lateral shift in the resulting array of images on the intermediate image plane (IIP). (B) Fabricated endoscope system. Scale bar: 100 mm. Red inset: Close-up of the working end of the endoscope, featuring a 7 GRIN lens array surrounded by illumination fibers. Scale bar: 5 mm.

**Figure 2.**
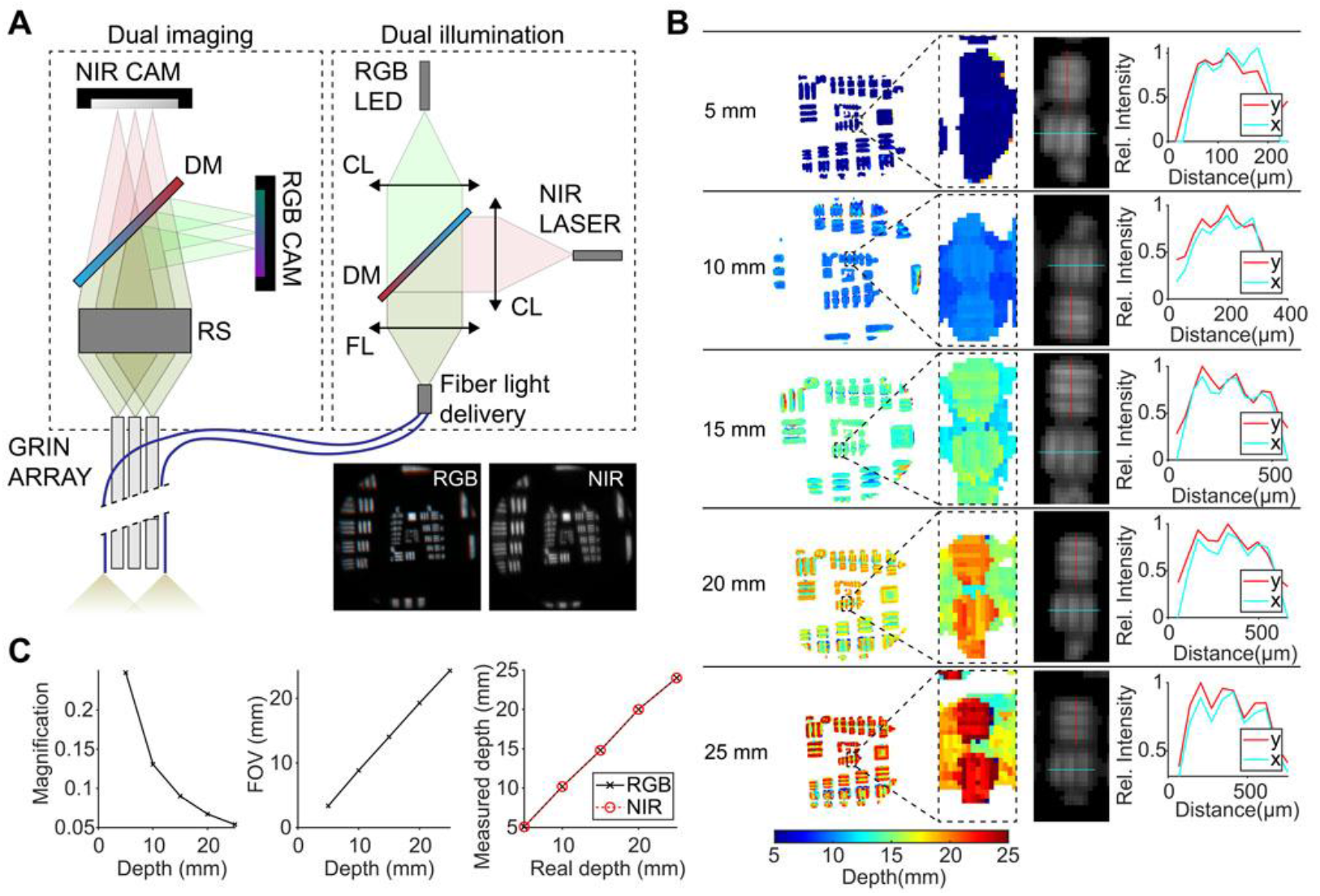
Endoscope layout and characterization. (A) Detailed schematic of optical component layout for dual imaging and illumination scheme. RS: relay stack. DM: dichroic mirror. CL: condenser lens. FL: focusing lens. (B) Characterization of system RGB imaging against USAF targets, extracting lateral resolution and depth. (C) Measured system characteristics of magnification (left), FOV (middle), and depth accuracy (right).

### 2.2 Flat target imaging

We first characterized the optical performance by calibrating the system’s PSF (Methods, Supplementary Note 1), then imaged a USAF target oriented normal to the optical axis at several depths in 5 mm increments. For each target depth, the light-field image was preprocessed and then reconstructed into a 3D volume using Richardson-Lucy Deconvolution (RLD). A depth map was extracted from each volume by finding the local maxima of intensity across depth for every lateral position (Supplementary Note 1).

As a result, we obtained depth maps that were well localized to the target’s actual depth (Figure 2B, C). Using the most in-focus slice of the volumetric reconstruction, we also characterized the lateral resolution of the reconstructed features (Figure 2B). Across the 20 mm axial range, our system was able to resolve bars with separations of 55.68 µm, 88.38 µm, 157.5 µm, 176.78 µm, and 198.42 µm in both directions (at 5, 10, 15, 20, and 25 mm distance from the endoscope tip, respectively). We next characterized the magnification over the entire imaging range, which decreased from 0.248× to 0.054× (Figure 2D). The FOV was calculated as the approximate diameter of the region where all 7 individual elemental views overlapped, and ranged from 3.4 mm to 24.3 mm over the entire imaging range (Figure 2E).

Similar results were observed on the NIR imaging channel. The depth accuracy of the NIR channel was the same as the color channel (Figure 2C), while the lateral resolution was slightly degraded along the x-axis. This effect was especially notable at a distance of 25 mm, exhibiting clear astigmatism (Figure S1) owing to the dichroic mirror’s placement within the converging path of light. However, the overall consistency of the resulting NIR volume indicates that it can be seamlessly integrated with the RGB volume.

### 2.3 3D object imaging

Next, we imaged a USAF target at a 45-degree tilt to better characterize the system’s 3D capabilities. As shown, the system clearly shows a smooth depth gradient along the tilted surface (Figure 3A, E). Our system was capable of distinguishing bars with axial separations down to 125 µm, although defocus artifacts elongate the axial extent of each feature, resulting in multiple local maxima along the depth of one bar feature (Figure 3E inset), and also result in spurious signal in the surrounding voxels (Figure 3A, E insets). As a result, the current depth localization strategy also assigns a depth for voxels outside of the known bar locations, causing ambiguity for smaller USAF elements (Group 2 elements 5 & 6), where each local peak in the depth profile did not necessarily correspond to a real bar (Figure 3B). Furthermore, for each 3-bar element, the axial distance between the uppermost and middle bar tended to be overestimated, while the distance between the middle and lowermost bar was underestimated (Figure 3C, G). We then computed the depth of each USAF element by taking the mean axial depth of each of the three bars. While the axial order between elements was maintained, quantitative distance measurements were sometimes underestimated by up to 25% when the target was positioned at greater distances (Figure 3D, H).

**Figure 3.**
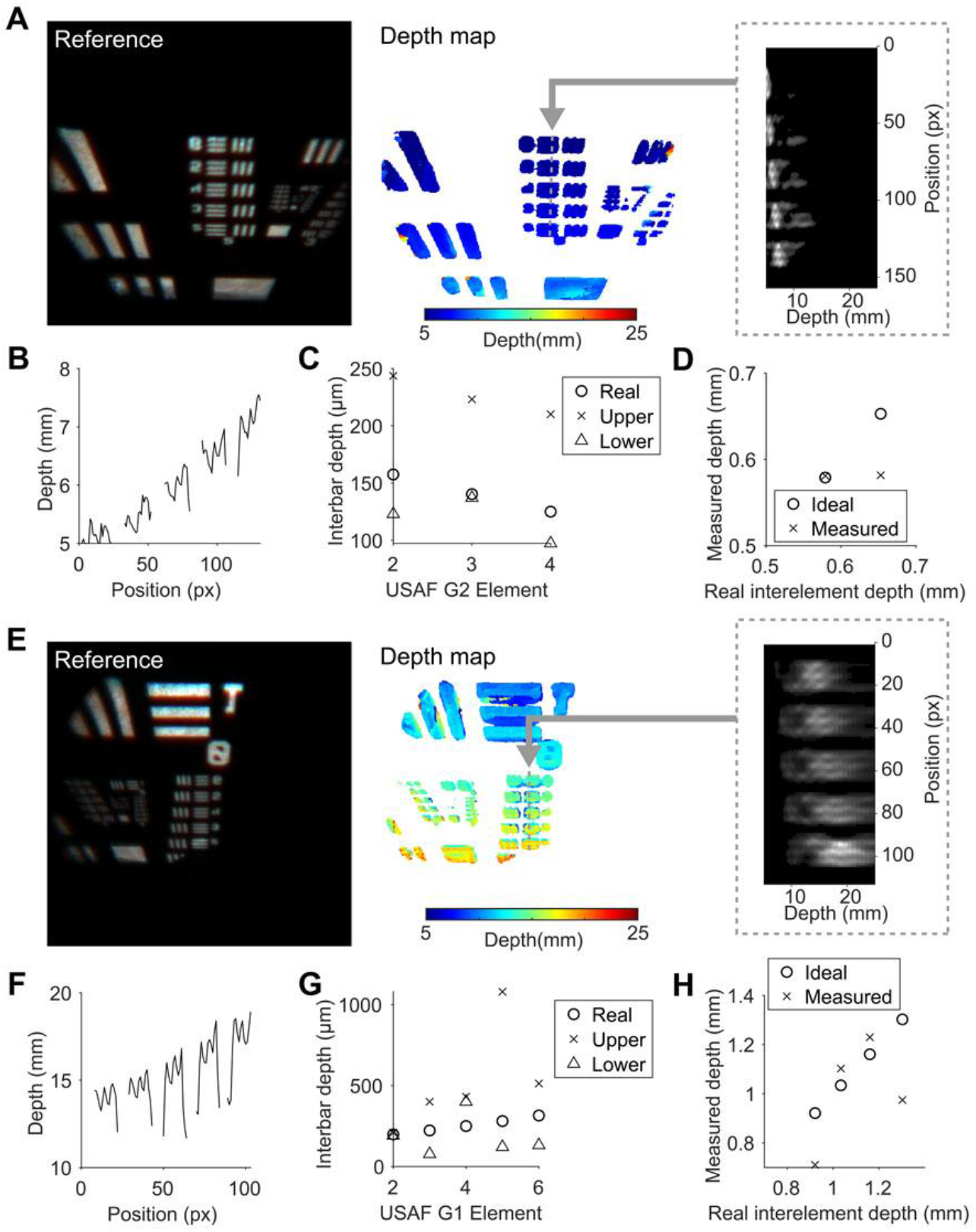
Tilted USAF target imaging. (A-D) Results corresponding to a tilted USAF target held at 5 mm away. (A) The central elemental image is provided as a 2D RGB reference, followed by the depth map calculated from the volumetric reconstruction. Dashed inset: The Z-Y projection of the volumetric reconstruction along the dotted grey line. (B) The surface depth along the dotted grey line. (C) Comparison of the depth between neighboring bars from the same USAF element. Circle: ground truth. X: Depth between the uppermost and middle bar of the group. Triangle: Depth between the middle and lowest bar of the group. (D) Comparison of the cumulative depth from the uppermost bar of USAF Group 2 Element 2 to the uppermost bar of other elements within the same group. (E-H) Results corresponding to a tilted USAF target held at 15 mm away. (E) The central elemental image is provided as a 2D RGB reference, followed by the depth map calculated from the volumetric reconstruction. Dashed inset: The Z-Y projection of the volumetric reconstruction along the dotted grey line. (F) The surface depth along the dotted grey line. (G) Comparison of the depth between neighboring bars from the same USAF element. Circle: ground truth. X: Depth between the uppermost and middle bar of the group. Triangle: Depth between the middle and lowest bar of the group. (H) Comparison of the cumulative depth from the uppermost bar of USAF Group 1 Element 2 to the uppermost bar of other elements within the same group.

To investigate the effectiveness of our system with different objects, we imaged various inorganic and organic targets (Figure 4). A paper “Georgia Tech” logo wrapped around a cylinder presented an accurate fitting to a circular surface profile with a 14.45 mm radius of curvature, the radius of the cylindrical substrate (Figure 4A). However, wrinkles on the back of the index finger knuckle resulted in a greatly exaggerated depth map. The larger wrinkles resulted in a depth >10 mm, compared to their actual depth of <1 mm relative to the surrounding tissue. Even smaller troughs result in a 3 mm depth change (Figure 4B).

**Figure 4.**
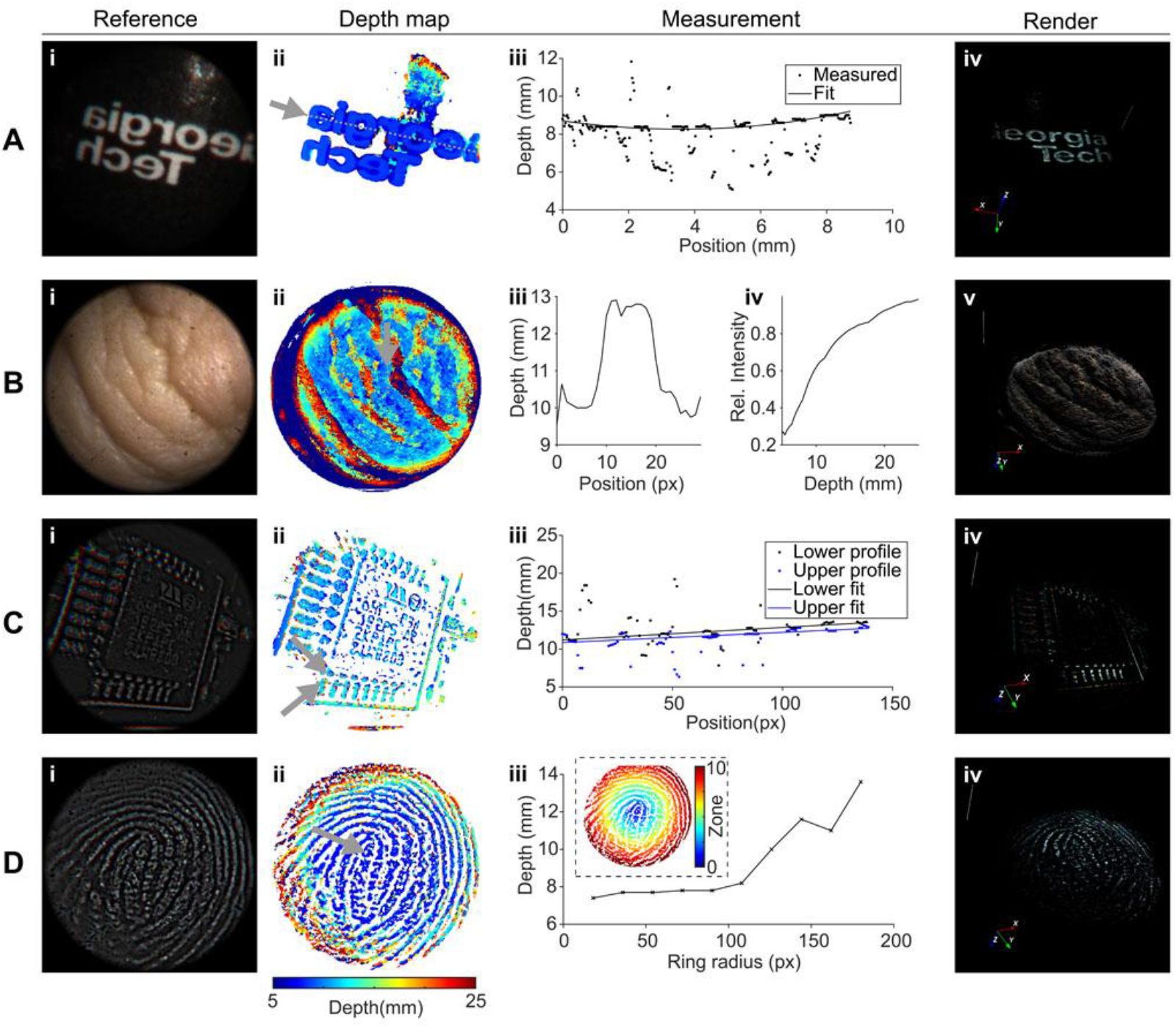
RGB volume reconstructions of various objects. (A) i. A “Georgia Tech” logo printed in white on a curved surface. ii. Reconstructed depth map. iii. Fitting the depth profile measured in A-ii (grey arrow and line). iv. Volumetric rendering of the object. (B) i. Skin wrinkles of the index finger knuckle. ii. Reconstructed depth map. iii. Depth profile measured across the profile in B-ii (grey arrow and line). iv. Mean intensity for each reconstruction volume slice. v. Volumetric rendering of the object. (C) i. Integrated circuit with contrast processing. ii. Reconstructed depth map. iii. Fitting a linear profile across the upper and lower parts of a series of soldered contacts in C-ii (grey arrow and line). iv. Volumetric rendering of the object. (D) i. Thumb fingerprint with contrast processing. ii. Reconstructed depth map. iii. Mean depth across a series of radial zones (defined in the dotted inset) extending from the center of the thumb. iv. Volumetric rendering of the object.

Thus, we find that Richardson Lucy Deconvolution faces challenges in volumetric reconstruction of opaque samples and in using maximum-intensity localization for 3D surface information (Supplementary Notes 2 & 3). To improve reconstruction quality for dense, opaque samples with a fixed degree of brightness, we introduced a feature enhancement scheme based on background subtraction (Methods, Supplementary Note 1). By subtracting a blurred version of the image from the original, we generate a sparse, high-contrast image with minimal background that highlights bright features. As a result, the reconstruction of complex objects improved greatly. For example, the contacts and even the imprinted text of an integrated circuit chip (Figure 4C) were reconstructed. By fitting a line profile across the upper and lower solder joints of a row of electrical contacts (Figure 4C-ii), we found the mean depth separation to be 0.557 mm, which differs by less than 0.5% from the expected true distance of 0.560 mm measured with a set of calipers (Figure 4C-iii).

We also measured the fingerprint texture of a thumb (Figure 4D). By taking the average depth across several radial zones, we characterize the curvature of the finger’s surface, showing a 0.8 mm drop from the center to a zone approximately 4 mm in radius. However, the average depth rapidly increases beyond this area, covering an additional 6 mm, which is not consistent with the imaged region, which has a total depth change of <2 mm. In part, this may be because, near the periphery of the image, not all elemental images intersect, leading to a loss of information. Furthermore, the structure density is higher than that of a computer chip.

### 2.4 Dual tissue imaging and biometric extraction

Finally, we unify the RGB and NIR speckle multimodal imaging capabilities of our system by computing 3D structure from both RGB images and flow maps. We first use a tilted microfluidic blood flow phantom, comprising square fluidic channels ranging from 300-1100 µm in width and height, perfused with an intralipid-based blood phantom (Supplementary Note 4). After processing the raw speckle elemental images into flow maps, we perform volumetric reconstruction on them to yield depth information consistent with the RGB channel reconstruction (Figure 5A, B)

**Figure 5.**
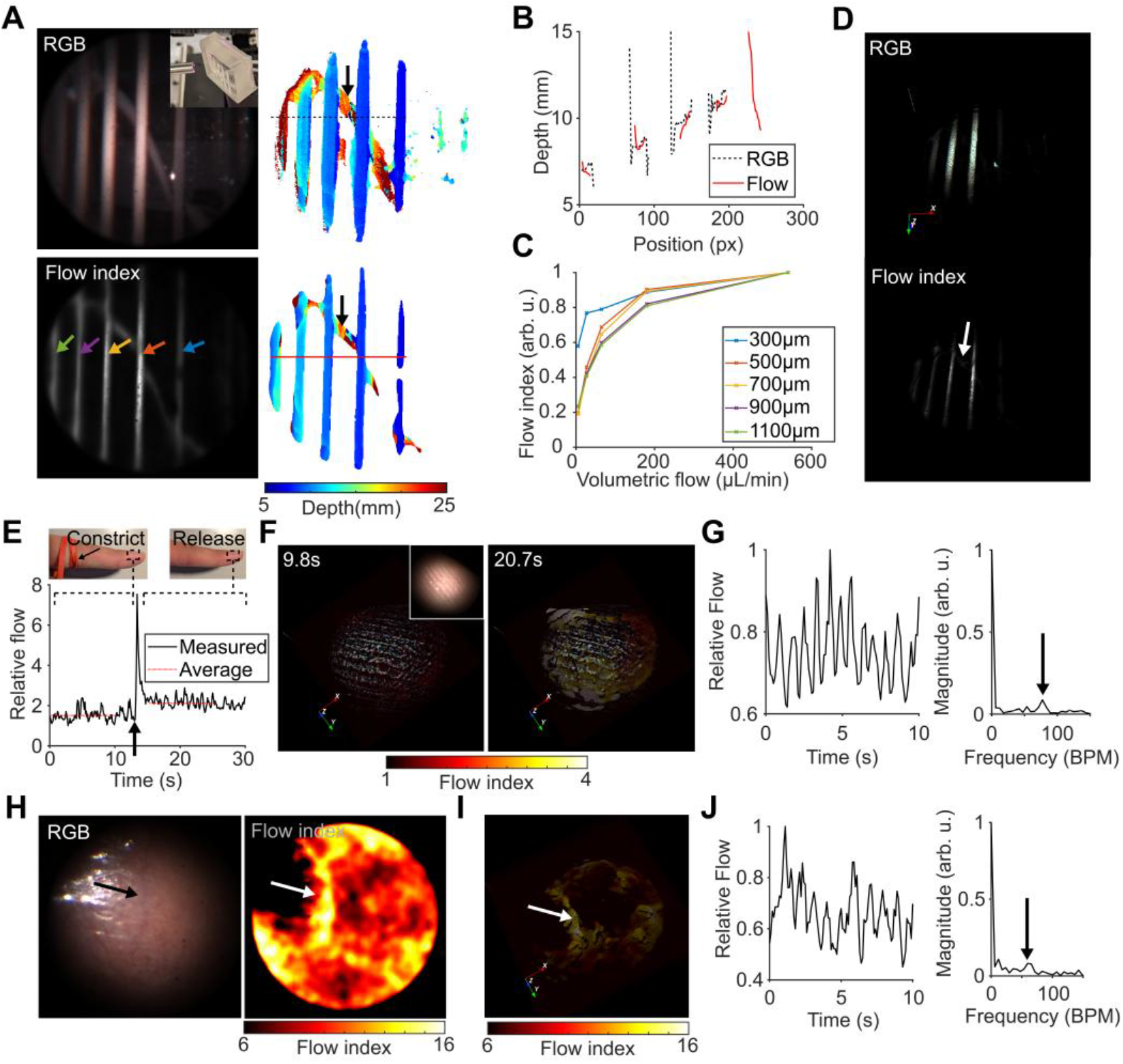
Multimodal imaging of flow phantoms and tissues under different physiological conditions. (A) Comparison of the RGB and NIR computed flow index central elemental image and their corresponding reconstructed depth maps. Black arrows: the liquid input tube located behind the phantom is accurately reconstructed. (B) Depths along the black dotted and red profiles for RGB and flow depth maps in (A). (C) Measured relative flow index compared to volumetric flow rate for each different channel size, indicated by the colored arrows in (A). (D) Volumetric rendering of each different information channel (RGB and Flow index). White arrow: The liquid input tube located behind the phantom is accurately displayed. (E) Measured the relative blood flow before and after the removal of a constricting finger blood flow tie. (F) Multimodal rendered volume showing the change in perfusion in a fingertip before and after the tie is removed. Flow information is presented as a colormapped overlay. (G) Measured relative flow of a finger at resting state with no tie, enabling the computation of pulse (right graph, black arrow). (H) Mucosa of the inner lip. NIR flow computation reveals a prominent blood vessel (white arrow), which is less visible in RGB (black arrow). (I) Render of the flow index, identifying the shallow vessel (white arrow). (J) Measured relative flow of the mucosa vessel at the resting state, enabling the computation of pulse (right graph, black arrow)

The system was perfused at volumetric flow rates of 5.4, 27, 64.8, 180, and 540 µL/min, which correspond to linear flow rates of 1-100 mm/s in the 300 µm channel, reflecting physiologically accurate blood flow rates in capillaries and arterioles [34–36]. Measuring the change in flow index across the different channels relative to the maximum volumetric flow rate of 540 µL/min, we observe that each shows a positive correlation. However, under ideal circumstances, this should be a linear relationship [32]. In this case, the imaged speckles are comprised entirely of subjective speckle, formed at the image sensor. While our previously reported proof-of-concept using an external single-mode fiber laser showed a highly linear trend [31], this new system uses multimode fibers for axial illumination, increasing the numerical aperture and enabling a larger light projection angle, thereby filling the FOV. However, multimode fibers support many propagating modes whose interference creates a static speckle pattern in the projected illumination, introducing a static speckle component that does not change with flow speed, thereby reducing the effective dynamic fraction (Figure 5C) [37]. As a result, our imaging conditions deviate from the assumptions required for simplified velocity computation and produce the observed asymptotic-like behavior. Our results indicate a key design consideration for further development: using single-mode fibers in conjunction with a compact beam-divergence scheme, such as an on-tip-fabricated lens [38].

Despite this, our current design’s relative perfusion measurement still enables us to monitor various biometrics. For example, we can detect changes in blood flow to the fingertip when a constricting tie is applied, which could be used to identify ischemic conditions (Figure 5E, F). Furthermore, larger vasculature is rendered visible with flow processing, enabling 3D localization of critical structures (Figure 5H, I). In both cases, we can use the temporal pattern of the flow index to identify the heart rate, indicating that our design can be used either to identify the general condition of a tissue or to target specific, larger individual vessels (Figure 5G, J).

## 3. Discussion

Our results demonstrate the feasibility of combining light-field with laser speckle contrast imaging in a single compact endoscopic device. Across visible and NIR imaging domains, the fabricated device achieves up to 55.68 µm lateral and 125 µm axial resolution at close range and has an imaging range of up to 20 mm with a maximum centimeter-scale FOV. Successfully integrating both imaging modalities yields both visual information and relative perfusion, enabling the simultaneous acquisition of both structural and biometric information.

However, our work also reveals several key considerations for the continued development of this technology. In particular, quantitative depth recovery was reliable only when sufficient contrast and sparsity were present in the image, performing well on synthetic targets, whereas biological tissues required contrast processing and faced additional challenges due to low texture. Our current approach relies on a maximum-intensity heuristic derived from the constructive signal of in-focus objects. While it works well on simpler, high-contrast objects, we observe that it is generally not robust to objects with low contrast or high background intensity. This weakness is shared with conventional depth-from-defocus methods relying on sharpness or gradients [39]. Indeed, tissues imaged during endoscopy are typically smooth and textureless, posing an ongoing challenge in 3D surgical imaging [40, 41].

Potentially, deep learning approaches have shown promise for light-field volumetric reconstruction. Not only can this approach improve reconstruction speed to near-real-time levels by eliminating the need for iterative processes [42, 43], but it has also been shown to overcome some challenges in the low-texture regime [44]. Furthermore, these networks can be augmented with physics-aware frameworks that take light-field properties into account [45], whilst large training sets can make reconstructions more robust to non-ideal conditions such as spatial variance, occlusion, or other conditions encountered in practice.

In summary, this work presents the first integration of light-field volumetric imaging with laser speckle perfusion mapping in an endoscopic form factor. The system successfully reconstructed 3D surfaces of high-contrast targets and detected relative perfusion changes in biological tissue. Looking forward, we anticipate that critical improvements in the robustness of depth reconstruction are the key research area to further enable this technology.

## 4. Methods

### 4.1. Endoscope design

Our system leverages a GRIN relay lens with a length >100 mm to achieve a long working length, resulting in considerable chromatic aberration (Figure S3). Consequently, we performed rigorous simulations to maximize imaging quality, decrease chromatic aberration, and maintain ease of fabrication by using commercial off-the-shelf (COTS) optics. A two-step MATLAB and ZEMAX optimization strategy was performed.

Because ZEMAX optimization cannot perform a discrete optimization over a catalog of lenses, we created a custom ray-optics-based genetic optimization scheme in MATLAB (Figure S3A) to generate a prototype design using a catalog of Thorlabs COTS lenses fitting general geometric and focus criteria, then followed up with fine-tuning of the optimized system in ZEMAX (Figure S3B). Based on paraxial ray optics principles, we programmed a ray-tracing simulation to determine the magnification and imaging plane at a given wavelength. Across a given lens with a focal length *f*, propagation from an object at a distance *d*_*o*_results in an image

*d*_*i*_ for a given wavelength *λ*:

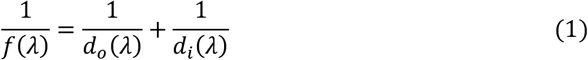

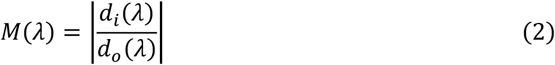

For singlet lenses, the wavelength-dependent focal length and principal planes were automatically calculated. Using 3-terms of Sellmeier coefficients for the lens material, we derived the refractive index *n* (*λ*) and applied it to the thick lensmaker’s equation to determine the nominal focal length of the lens, *f* (*λ*), and principal plane locations *h*_1_(*λ*), *h*_2_(*λ*):

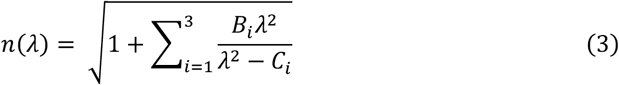

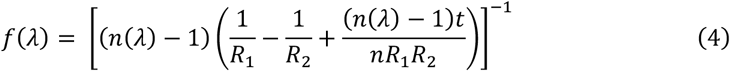

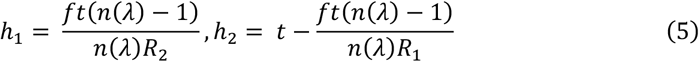

Singlet lens radii *R*_1_, *R*_2_, thickness *t*, and Sellmeier coefficients *B*_*i*_, *C*_*i*_ were obtained from the manufacturer’s catalog. For achromatic doublets, these properties were manually exported from the manufacturer’s provided ZEMAX files.

Thus, the propagation of an object across the *i*th lens located can be modeled as:

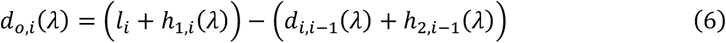

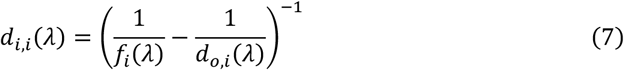

where *l*_*i*_ is the distance between the first surface of the *i-1*th lens and the *i*th lens, *h*_1,*i*_, *d*_*o,i*_ are with respect to the first surface of the *i*th lens, and *h*_2,*i*−1_, *d*_*i,i*−1_, are with respect to the first surface of the *i-1*th lens. In the case of propagation across the first lens, *d*_*i*,0_ = 0 and *h*_2,*i*−1_ = 0.

To limit the search space, our system was restricted to two convex and one concave lens, and two system configurations were allowed: the concave lens placed at L2 or at L3 (Figure S3A). Six additional variables were optimized over: the type model of each of the three lenses (L1, L2, L3), which were drawn from a discretized catalog of convex or concave lenses. The distance between the first lens and the nominal intermediate image plane (d1). The distance between the last two lenses (d2). And, the magnification. These variables would be enough to constrain the overall system. Additional boundary conditions were implemented to limit the magnification factor and ensure the system lengths remain positive. The merit function was to minimize the separation *S* between the two most extreme wavelength-dependent image planes:

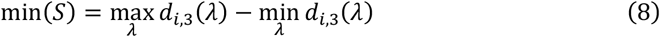

The MATLAB genetic algorithm solver was used to optimize these seven variables with an initial population of 100,000. The prototype solution was then implemented into ZEMAX for further fine-tuning. The performance of the optimized simulation showed a clear decrease in chromatic aberration, maintaining a more equal intensity ratio among the R, B, and G channels in the imaging spot across a wide field angle and depth (Figure S4). However, a minor degradation in the FWHM was observed in the system (Figure S4I-J).

The as-built system matches the performance observed in the simulations, and we observed a clear decrease in chromatic aberration, allowing us to resolve the gaps in the blue-channel targets (Figure S3C-F). However, it does trade off some resolution in the red and green imaging channels.

### 4.2. Endoscope instrumentation

The endoscope was designed with a blend of optical cage components and custom 3D-printed housings (Form 2, Formlabs), and aimed to maximize the ease of assembly, disassembly, and calibration adjustments (Figure S5). Seven 1 mm diameter GRIN lenses (GT-ERLS-100-005-250, GRINTECH) were slotted into a 3D-printed working-end optics holder. A 1:7 fiber optic fan-out (BF72HS01, Thorlabs) was used as the illumination. The ferrule of six of the seven fiber optic outputs was removed via heat treatment, and the fiber ends were then slotted into the working-end optics holder. Afterwards, working end of the endoscope was sheathed using a gauge 6RW stainless steel tube (New England Small Tube Corporation), and sealed using a glass window (22-036, Edmund Optic) and optical adhesive (Norland 68, Edmund Optics), rendering the working end waterproof.

A custom 3D-printed camera housing was fabricated. A 750 nm cutoff longpass dichroic mirror (69-882, Edmund Optics) was inserted at a 45-degree angle into the camera housing. A monochrome (acA4024-29um, Basler) and RGB (acA4024-29uc, Basler) camera were assembled on the transmissive and reflective paths of the dichroic mirror, respectively. An additional 700 nm cutoff long-pass filter (15-228, Edmund Optics) was placed in front of the monochrome camera to further restrict light to the NIR range.

The relay optics stack was assembled in standard 1” lens tubes and consisted of a 60 mm achromat (AC254-060-A-ML, Thorlabs), followed by a 3D-printed 5.2 mm diameter aperture, a 40 mm achromat (AC254-040-A-ML), and finally a −75 mm biconcave lens (LD1170-A, Thorlabs).

The working end, lens stack, and main body of the endoscope were then assembled together. Alignment was performed by imaging a USAF target at 7 mm, while the position of the individual GRIN lenses was manually adjusted. Once aligned, the lens positions are fixed by depositing glue into specialized access slots.

The dual light source was assembled using an LED-coupled white lamp (MWWHF2, Thorlabs) with a 400 µm core multimode fiber. The lamp was collimated with a f = 16 mm condenser lens (ACL25416U-A, Thorlabs), and the beam passed through a 750 nm cutoff short-pass dichroic mirror (69-219, Edmund optics) oriented at a 45-degree angle. The NIR branch of the illumination comprises a 785 nm laser diode (L785P090, Thorlabs), which was corrected for ellipticity using a f=75 mm plano-convex cylindrical lens (LJ1703RM, Thorlabs) and collimated by a f=75 mm lens (AC254-075-B-ML, Thorlabs). The diode was controlled using a laser diode controller (TLD001, Thorlabs) and operated at 135 mA. The two beams were combined with the dichroic mirror, then focused by a f=30 mm lens (AC254-030-AB-ML, Thorlabs) onto the face of the 1:7 fiber fan-out bundle. A piece of scotch tape was placed on the fiber input surface to diffuse the focus spot, making it more uniform and improving the fiber light-filling conditions.

### 4.3. Instrument calibration

Briefly, a distortion and magnification calibration was performed by imaging checkerboard targets. Each elemental view was cropped from the raw camera sensor image, then corrected for lens distortion using OpenCV. Magnification across all elemental views was normalized using linear interpolation.

Next, a one-time PSF acquisition was performed by imaging a back-illuminated 10 µm pinhole across a 3-28.5 mm axial range with a step size of 25 µm. The experimental PSF was corrected by normalizing the spot energy, distortion, and magnification of each elemental image. The experimental PSF was then converted into a hybrid PSF by replacing the centroid of each elemental spot with a Gaussian kernel to improve reconstruction resolution. Finally, the PSF was axially downsampled using a geometric downsampling scheme, in which the depth step size increased from 25 µm closest to the endoscope to 800 µm at the furthest axial range, reducing the PSF to a total of 128 axial slices spanning the depth.

### 4.4 Laser speckle processing

Raw speckle elemental images were first corrected for distortion and magnification. For every pixel, the speckle contrast *K* was calculated [46]:

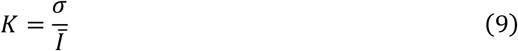

where *σ* is the intensity standard deviation over some spatio-temporal pixel window, and *Ī* is the intensity mean of the same window. In our experiments, we utilize a 5-by-5-by-3 spatiotemporal window, yielding elemental contrast maps *K*_*e*_. Owing to the stochastic nature of the speckle process, the flow index images appeared grainy. A guided filter *G* [47] was applied, using the greyscaled RGB image as the reference, to smooth the flow index while retaining structural information.

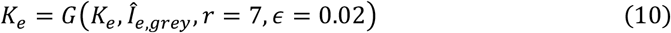

The flow index *f* is proportional to the underlying flow speed *v* and is approximated as [32].

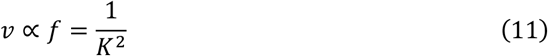

To prevent infinite values resulting from dark areas with no signal, the flow index was capped at 200, which was empirically determined to be a reasonable maximum value that real media could not exceed. The flow index images *f*_*e*_ are subsequently treated as normal light-field images in the rest of the pipeline and reconstructed using RLD. For a single light-field image, speckle processing took 50 ms.

### 4.5. Volumetric reconstruction

A custom GPU-accelerated implementation of 3D RLD was implemented using the CuPy package in Python. A depth regularization scheme was implemented to counteract observed intensity accumulation along the depth of the reconstruction.

Prior to volumetric reconstruction, raw images were preprocessed to correct for distortion and magnification (Supplementary Note 1). Depending on the sample, feature enhancement was applied to low-contrast images (biological samples, objects) by subtracting a non-local means (NLM) filtered version of the image (Supplementary Note 1). Each image channel (R, G, B, flow) was processed independently for 30 iterations of RLD, which took approximately 10 seconds on our current hardware (NVIDIA RTX 3060). The resulting volume was centrally cropped to match the size of the central elemental view, and the top and bottom 16 depth layers were cropped, yielding a final volume of 396 × 396 × 96 voxels. For RGB volumes, the volumes for each color channel were then subsequently concatenated.

To extract depths from the volumetric data, each depth in the deconvolved volume was multiplied by the corrected central elemental image to mitigate defocus artifacts, and then the resulting image was thresholded. The true feature depth at a given lateral position was then defined as the location of the first local intensity maximum along the z-axis, and this value was extracted. The final volume was rendered using the VTK library [48].

### 4.6 Imaging

#### Optical targets

The system was characterized using a negative USAF target (R1DS1N, Thorlabs) illuminated using a collimated white LED (Thorlabs MMF2). A diffuser was placed immediately behind the target. The target was stepped between 5 and 25 mm from the endoscope and tilted to 45 degrees for angled imaging trials.

#### Miscellaneous objects

A black-and-white Georgia Tech logo was printed on paper and then taped onto a cylindrical beaker with a diameter of 28.9 mm. After imaging with the endoscope and reconstruction without contrast filtering, depth across a linear profile was extracted from the depthmap using ImageJ, then fit to a circular function with a radius of 14.45 mm using MATLAB:

The back of the index finger knuckle, with the finger outstretched, was imaged using the system and then volumetrically reconstructed without contrast filtering. Depth along a linear profile was extracted using ImageJ.

The integrated circuit was imaged and volumetrically reconstructed with contrast filtering. The normal of the chip surface was determined by fitting a line to each perpendicular set of contacts along one axis of the chip, then computing the cross-product of the direction vectors of these lines. Accordingly, the expected separation (0.56 mm) between the lower and upper joints of the contacts along the optical axis was determined by projecting the distance measured with calipers (0.6 mm) along this vector.

The central whorl of the thumb was imaged and volumetrically reconstructed with contrast filtering. Because of the many edges and features of the fingerprint, the output surface map had a noisy texture. Thus, rings were defined with 18-pixel radius increments, and the mean depth within each ring was calculated, revealing the gradual curvature of the center finger.

#### Blood phantom

A scattering-matched blood phantom was flowed through a custom microfluidic phantom with square channels ranging between 0.1 – 1.1 mm in cross-sectional side length (Supplementary Note 4). For each experiment, a reference RGB image of the microfluidic chip with no flow was first captured using the endoscope. Then, a syringe pump (Pump 11 Elite Infuse Only, Harvard Instruments) was used to infuse the microfluidic chip at 5 different volumetric flow rates of 540, 180, 64.8, 27, and 5.4 µL/min. For each flow rate, infusion was performed for two minutes, then paused for 2 minutes. Laser speckle video was captured during the infusion experiment, with NIR laser illumination on, at 10 FPS for 20 minutes with a 10ms exposure time. Afterward, raw light-field speckle images were processed to compute the flow index and then reconstructed alongside the RGB reference image.

#### Biological sample imaging

To image the finger under different flow conditions, the index finger was held on a flat, stable platform, and a rubber band was wrapped tightly around its base. After several seconds of recording, the rubber band was released, allowing reperfusion of the finger.

To record the inner lip mucosa, a volunteer held their head steady on a flat surface and used their finger to invert the lower lip. Several seconds were recorded from the endoscope, during which the volunteer was allowed to breathe normally.

Trials consisted of capturing an RGB reference image, followed by capturing a laser speckle video at 10 FPS with a 10 ms exposure. After video capture, raw light-field speckle images were processed to compute the flow index and then reconstructed alongside the RGB reference image. Owing to the smooth, textureless surface of the inner lip mucosa, the RGB volumetric reconstruction failed to produce an accurate representation of the surface and was omitted from the scene rendering.

To measure biometric signals, the flow index from the central elemental view was used. For total finger perfusion, the mean flow index for every frame was calculated. The Fourier transform of the resulting time signal, along with the non-DC local maxima, identified the pulse rate. For the inner lip mucosa vessel, basic motion correction was performed to compensate for natural motions such as breathing.

## Acknowledgment

We acknowledge the support of the NSF-GRFP, NIH, and NSF programs.

## Data availability statement

Data underlying the results presented in this paper are not publicly available at this time but may be obtained from the authors upon reasonable request.

## Supplementary Document

See Supplement 1 for supporting content.

