## Supplemental document 1 for "Multimodal 3D light-field and laser-speckle endoscopy"

### Multimodal 3D light-field and laser-speckle endoscopy: supplemental document


#### Note S1: Detailed processing pipeline

##### *Distortion and magnification calibration*

First, the centers of each elemental view were found. Using the white reference image, image binarization was performed. The diameter of the elemental views on the camera was manually measured at $L = 792$ pixels. Then, circles of the same diameter were detected using the Hough transform, resulting in the identification of the center coordinates $c_e$ of each of the elemental images. Thus, to extract the elementals from any raw light field image, square areas centered on each were cropped out.

To ensure the validity of the shift-invariant assumption required for Richardson-Lucy Deconvolution (RLD), a two-stage geometric calibration was performed for each elemental view $e$.

First, we captured at least 10 images of a chessboard target for each perspective. These were used to compute a unique distortion mapping, $\mathcal{D}_e$, for each elemental view using the OpenCV framework [1] (**Figure S7**A).

Next, the average relative magnification factor $\mathcal{M}_e$ between a peripheral elemental view and the center elemental view was determined. This step is critical, as RLD relies on the assumption of a laterally-shift invariant PSF at each depth. However, magnification differences among elemental views guarantee that this assumption is false and will therefore cause reconstruction errors. While we cannot fully correct for magnification differences, as they arise from variations in focal length and lens position and therefore vary across the imaging depth, we can reduce their impact by applying a constant magnification correction to each elemental view. A printed grid target was translated axially from 3 - 28.5 mm with a 25 µm step size. At each step, the length of the imaged grid squares in each peripheral view $I_e(z)$ was measured relative to the central view $I_{e_{central}}(z)$ (**Figure S7**A). The resulting relative magnification curves (**Figure S7**B) were fit to asymptotic functions. We averaged the relative magnification factor over the 5–15 mm range—the region of highest variance—to yield a fixed factor $\mathcal{M}_e$ for each elemental view.

Future elemental images are corrected by undistorting and then rescaling the image by the magnification factor using a bilinear-interpolation-based image resizing function $\mathcal{B}$ (Eq. $\mathrm{S}-1$). Applying these corrective factors greatly improved the quality of the volumetric reconstructions (**Figure S7**C).

$$\hat{\boldsymbol{I}}_e = \boldsymbol{\mathcal{B}}\left(\boldsymbol{\mathcal{D}}_e{}^{-1}(\boldsymbol{I}_e), \boldsymbol{\mathcal{M}}_e\right) \qquad (\mathrm{S}-1)$$

*PSF collection*

The PSF collection and calibration were performed by imaging a 10 μm pinhole back-illuminated by a fiber-coupled broadband LED (MWWHF2). Notably, the LED emits light in the 700 -800 nm range. A piece of translucent tape serving as a diffuser was applied to the back of the pinhole, and light was then collimated and focused onto the pinhole, which was mounted on a motorized stage (NRT150, Thorlabs). The imaged pinhole was centered within the central elemental image. Then, an automated multi-exposure PSF acquisition was performed. Using a 25 μm step size, the pinhole was moved from 3 mm to 28.5 mm from the endoscope, resulting in 1021 depth steps. At each step, exposure images of 1, 4, 8, and 16 ms for RGB and 4, 40, 160, and 400 ms for the NIR camera were taken, ensuring that both the center and peripheral elemental images captured the PSF spots with good dynamic range across the entire imaging depth. Finally, a dark image with no illumination and a white image in which collimated light was shone directly onto the endoscope were taken.

*PSF calibration*

Next, for a given depth, each exposure image was read. The elemental views were extracted and assembled into a tensor $PSF_{meas,z}$ of size of [*E, P, C, L, L*], where *E* is the number of elemental views (7), *Q* is the number of exposures, *C* is the number of color channels (3 for RGB, 1 for NIR), and *L* is the width and height of the elemental image (792 pixels). For every elemental view $e \in E,$ and for every color channel $c \in C$, one exposure $\hat{q}$ was selected that would maximize the intensity value of the image without saturation of the 8-bit dynamic range. This process was repeated for each depth $z \in Z$, resulting in a multidimensional array $PSF_{raw}$ of data of the size [*E, Z, C, L, L*], where *Z* is the total number of depth steps (1021):

$$\boldsymbol{PSF_{raw}(e,z,c,i,j) = \ PSF_{meas,z}(e,\hat{q},c,i,j)} \qquad (\mathrm{S}-2)$$

$$\boldsymbol{\hat{q} = \underset{q \in Q}{\mathrm{argmax}} \left\{ q \middle| \max_{i,j \in L} PSF_{meas,z}(e,x,c,i,j) < 255 \right\}} \qquad (\mathrm{S}-3)$$

Then, a spot energy normalization was performed in which each elemental image was divided by the sum of intensity within a 128-by-128 box centered about the brightest pixel, represented by Ω:

$$\boldsymbol{PSF_{norm}(e,z,c,i,j) = \frac{PSF_{raw}(e,z,c,i,j)}{\sum_{(i,j)\in\Omega} PSF_{raw}(e,z,c,i,j)}} \qquad (\mathrm{S}-4)$$

The entire calibration process was repeated for each PSF type (RGB or NIR).

*Formation of hybrid or real PSFs*

We experimented with two types of PSFs: the experimental PSF and the hybrid PSF, which replaces the experimental spot with a Gaussian spot at the same location. For every elemental view *e*, depth *z*, and color channel *c*, the centroid coordinate $p$ of the PSF spot was calculated. The drift of the central elemental spot across the depth was first calculated and used as an offset $\Delta p$ for all centroids, correcting misalignment of the ideal on-axis PSF spot.

$$\boldsymbol{\Delta p(z,c) = p(e_{center},c,z) - \ p(e_{center},c,z_0)} \qquad (\mathrm{S}-5)$$

Distortion and magnification correction were then applied to the resulting coordinates:

$$\boldsymbol{\hat{p}(e,c,z) = \mathcal{M}_e \cdot \mathcal{D}_e^{\ -1}\big(p(e,c,z) - \Delta p(z,c)\big)} \qquad (\mathrm{S}-6)$$

Finally, each PSF image was resampled via 2D interpolation to place the experimental spot at the ideal corrected centroid coordinate $\hat{p}$, yielding the corrected experimental PSF (**Figure S8**A).

To form the hybrid PSF, we initialized a null tensor of dimension [*E, Z, C, L, L*] and placed a Gaussian spot at the location of the corrected centroid coordinates generated during the experimental PSF processing. The size of the Gaussian kernel was matched to the experimental data by approximating the full width at half maximum (FWHM). For an experimental PSF spot at *e, c,* and *z*, we sum the number of pixels that exceed half the maximum image intensity, then calculate the equivalent radius $r$.

$$\boldsymbol{r(e,c,z)} = \sqrt{\frac{1}{\pi}\sum_{\boldsymbol{i,j}}\left[\boldsymbol{P_{norm}(e,c,z,i,j)} > \boldsymbol{0.5}\max_{\boldsymbol{i,j}}\boldsymbol{P_{norm}(e,c,z,i,j)}\right]} \qquad (\mathrm{S}-7)$$

The Gaussian spot is then approximated as

$$\boldsymbol{\sigma(e,c,z)} = \frac{\boldsymbol{r(e,c,z)}}{\boldsymbol{2.355m}} \qquad (\mathrm{S}-8)$$

where *m* is a contraction factor introduced to modify the size of the Gaussian kernel. The contraction factor of 3 was chosen to provide the best reconstruction quality while minimizing modifications to the measured PSFs (**Figure S8**B, E-G). This process was repeated for each PSF type (RGB or NIR).

Subsequently, a nonlinear PSF depth-sampling strategy was employed to improve computational efficiency. Using a full-resolution PSF is too computationally infeasible, and furthermore, the lateral shift of the PSF centroid per axial step diminishes as the distance from the endoscope increases (**Figure S8**C). Consequently, uniform axial sampling at 25 µm results in oversampling at greater depths. To address this, we employ geometrically-strided resampling, extracting the PSF data along *Z* starting every n = 1 (25 µm) slices, doubling along the depth range up to n = 32 (800 µm) slices, the furthest depths (**Figure S8**D). To minimize intensity accumulation artifacts at the volume boundaries, we sampled 16 buffer slices at each end of the PSF with fine depth increments, which are cropped from the final reconstructed volume. We achieve an approximately 8× data compression in $PSF_{reduced}$, reducing the $Z$ dimension to 128 elements

To prepare for use in RLD, a light field mosaic of $PSF_{reduced}$ is constructed by closely tiling the elemental views in the lateral plane, thereby reducing the void space in the raw light field image. The mosaic light field image was downscaled by a factor of 2 to allow efficient RLD computation. To avoid boundary wrapping during the Fourier transform, the mosaic was zero-padded to a 2048-by-2048 lateral dimension, yielding the final light-field PSF = $PSF_f(c,z,i,j)$.

##### *Preprocessing for RLD*

Prior to performing RLD, raw light field images are preprocessed by extracting the elemental views $I_e$ and applying distortion and magnification correction as described previously. Depending on the contrast of the image (**Figure 4**B), we also applied a feature enhancement approach by subtracting a non-local means (NLM) filtered version of the image from itself[2]:

$$\widehat{\boldsymbol{\mathcal{L}}}_{\boldsymbol{f}} = \boldsymbol{\mathcal{L}_f} - \mathbf{NLM}(\boldsymbol{\mathcal{L}_f}) \qquad (S-9)$$

This serves to remove a constant-intensity background and enhance feature contrast, thereby reducing depth-biasing during RLD reconstruction.

Then, the elemental images are assembled into a compact light-field mosaic by closely tiling them. The same mosaicking process is applied to the hybrid PSF, saving memory by reducing the void space between elemental views compared with the raw light-field image. The resulting mosaics are downscaled by a factor of 2 to allow efficient RLD computation. To avoid boundary wrapping during the Fourier transform, each mosaic was zero-padded to a 2048-by-2048 lateral dimension, forming the final light-field PSF $PSF_f(c, z, i, j)$ and final light-field mosaic $\mathcal{L}_f$.

##### *Volumetric deconvolution*

A custom GPU implementation of 3D Richardson-Lucy Deconvolution (RLD) was written in Python, leveraging the CuPy libraries[3]. The standard RLD algorithm is described as:

$$\boldsymbol{v}_{k+1} = \boldsymbol{v}_k \cdot \boldsymbol{\mathcal{H}}^T * \left(\frac{\widehat{\boldsymbol{\mathcal{L}}}_f}{\boldsymbol{\mathcal{H}} * \boldsymbol{v}_k}\right) \qquad (\mathrm{S}-10)$$

Adaptation to the 3D case requires modifications to the forward and backward projections [4-6]. The forward projection $\mathcal{H} * v_k$ mapping the volume to an estimated camera image is a summation of layer-by-layer convolutions between the 3D $PSF_f$ and volume $v_k$ across *z*.

$$\boldsymbol{\mathcal{H}} * \boldsymbol{v}_k = \sum_{\boldsymbol{z}} \mathbf{PSF_f}(\boldsymbol{z}) * \boldsymbol{v}_k(\boldsymbol{z}) \qquad (\mathrm{S}-11)$$

Conversely, the backward projection $\mathcal{H}^T * \mathcal{R}$ is a convolution between the rotated PSF and the residual $\mathcal{R}$, broadcast over all *z* of the rotated PSF.

$$\boldsymbol{\mathcal{H}}^T * \boldsymbol{\mathcal{R}} = \underset{\mathbf{180}}{\mathbf{Rot}}(\mathbf{PSF_f})(\boldsymbol{z}) * \left(\frac{\widehat{\boldsymbol{\mathcal{L}}}_f}{\boldsymbol{\mathcal{H}} * \boldsymbol{v}_k}\right) \qquad (\mathrm{S}-12)$$

Due to the observed accumulation of intensity along the depth of the reconstruction (**Figure S2**), we also implemented a depth regularization scheme during deconvolution. At each iteration of the deconvolution, the projected volume intensity is decayed by a factor of $\mathrm{r(z)} = 1 - 0.002 i_z$, where $i_z$ is the index corresponding to the depth of a slice in the volume:

$$\boldsymbol{v}_{k+1} = \boldsymbol{v}_k \cdot \boldsymbol{\mathcal{H}}^T * \left(\frac{\widehat{\boldsymbol{\mathcal{L}}}_f}{\boldsymbol{\mathcal{H}} * \boldsymbol{v}_k}\right) * \boldsymbol{r} \qquad (\mathrm{S}-13)$$

Due to the large image sizes, high consideration was given to memory management. Towards this end, for RGB image inputs, $PSF_f$ was averaged across all color channels. The PSF and its rotation were stored in GPU memory after the Fourier transformation, reducing the number of Fourier transform operations. The most memory-efficient implementation of RLD while still maintaining reconstruction speed requires 3.5× the memory of the optical transfer function ($OTF = \mathcal{F}(PSF)$), which keeps all values on-GPU and avoids expensive memory transfers. For a single channel, 30 reconstruction iterations took 10s on our hardware, which could be improved through patch-wise approaches[7].

The deconvolved volume has a dimension of 2048-by-2048-by-128. 16 buffer depths from the front and end of the volume are removed, and the central portion of the volume matching the size of the central elemental view is cropped, resulting in a final volume of 396-by-396-by-96 voxels. For RGB input images, each color channel was reconstructed independently, then merged to result in the final color volume.

##### *Surface finding*

Although RLD provides a volumetric reconstruction of the data, it can only estimate depth, as objects still exhibit defocusing patterns at other z-planes. Thus, we devised a straightforward depth-finding method by determining the maximum-intensity depth at each lateral position.

First, we devised an image processing scheme to reduce the impact of defocus artifacts. Each depth of the reconstructed volume $\mathcal{V}$ was multiplied by the corrected central elemental image, thereby reducing defocus artifacts that appear between features.

$$\boldsymbol{\mathcal{V}'(x,y,z,c) = \hat{I}_{central}(x,y,c) * \mathcal{V}(x,y,z,c)} \quad \text{(S} - 14\text{)}$$

Then, a threshold operation was performed to eliminate spontaneous low-intensity pixels that do not correspond to any real features.

$$\boldsymbol{\mathcal{V}_{filter}(x,y,z,c) = \begin{Bmatrix} \mathcal{V}'^{(x,y,z,c)} & \text{if } \mathcal{V}'(x,y,z,c) \geq T \\ 0 & \text{if } \mathcal{V}'(x,y,z,c) < T \end{Bmatrix}} \quad \text{(S} - 15\text{)}$$

Finally, the intensity profile $Q$ along z, averaged across all channels, was extracted, then thresholded to values above 75% of the maximum profile intensity.

$$\boldsymbol{Q_{i,j}(z) = \overline{V}_{filter}(i,j,z)} \quad \text{(S} - 16\text{)}$$

$$\boldsymbol{Q'_{i,j}(z) = \begin{Bmatrix} Q_{i,j}(z) & \text{if } Q_{i,j}(z) \geq 0.75 \max_z Q_{i,j}(z) \\ 0 & \text{if } Q_{i,j}(z) < 0.75 \max_z Q_{i,j}(z) \end{Bmatrix}} \quad \text{(S} - 17\text{)}$$

The depth map $\hat{z}$ was then estimated as the $z$ of the first occurring local maxima. This method prevents bright artefacts at deeper layers from overriding the depth measurement, as the reconstruction process is known to have an intensity bias towards the deeper volume areas (**Figure 4B**-iv).

##### *Visualization*

The VTK library [8] was used to visualize the 3D voxel volume. First, the depth map was utilized to refine the volume. At each lateral position, only the voxel corresponding to the mapped depth was retained while all others along $z$ were set to 0.

$$\boldsymbol{\mathcal{V}_{refine}(x,y,z,c) = \begin{Bmatrix} \mathcal{V}(x,y,z,c) & \text{if } z = \hat{z}(x,y) \\ 0 & \text{if } z \neq \hat{z}(x,y) \end{Bmatrix}} \quad \text{(S} - 18\text{)}$$

A custom render mapping was written to visualize the resultant volume, showing the RGB volume voxel values as RGBA, where the alpha (A) channel was computed according to mean brightness over all colors.

For flow visualization, we instead used a different approach. Because the RLD process is nonlinear, the relative flow index information would be lost after reconstruction. Thus, the initial RLD volume reconstructed from the flow maps was utilized to derive the volume depth map, and the flow magnitude of the 2D central elemental image was mapped onto this surface:

$$\boldsymbol{\mathcal{V}_{flow}(x,y,z) = \begin{Bmatrix} f_{central}(x,y) & \text{if } z = \hat{z}(x,y) \\ 0 & \text{if } z \neq \hat{z}(x,y) \end{Bmatrix}} \quad \text{(S} - 19\text{)}$$

To combine the two views, a colormap was applied to the flow volume, and the RGB volume and flow volume were balanced at 50% transparency each.

##### Note S2: Discussion on depth extraction and its limitations.

Localization of each feature's depth was performed by finding the local intensity maxima along the depth for each lateral position. This heuristic is motivated by the underlying mathematical properties of the RLD algorithm. During the backprojection step, the residuals are convolved with the flipped PSF to result in the estimated volume. If the residual is initialized as the light-field image in the first iteration, this is analogous to a shift-and-sum operation of the elemental views [9]. While RLD differs in its application of the wave-optics PSF model and in the continued refinement of the estimated volume via forward- and back-projection iterations, conceptually, the in-focus signal from features at a given depth should constructively align, and the out-of-focus signal should be blurred.

However, our results show that this process is not robust for opaque samples such as tissues. We rely on the constructive addition of brightness to form features; however, this can also cause mischaracterization of depth due to defocusing. At off-depth planes, virtual defocus can cause light from a bright feature to bleed into neighboring pixels corresponding to relatively darker features, thereby introducing crosstalk in the depth calculation based on the reconstructed defocus pattern. Thus, a certain degree of spatial sparsity is required to enable more accurate localization. Accordingly, we applied a feature enhancement scheme by subtracting a spatially blurred version of the image from itself to improve the contrast of key features (Supplementary Note 3).

As noted in the Discussion, this weakness is shared with other 3D imaging approaches [10] and is further complicated by the smooth, relatively low-texture tissues encountered during endoscopy [11, 12]. Looking towards the future, major advancements will focus on improving the applicability of our 3D endoscopic system to these surgical imaging conditions, potentially through deep learning approaches [13, 14].

#### Note S3: Discussion on Richardson Lucy depth regularization.

Typically, light field imaging is performed in fluorescence microscopy, where the imaged volume is largely transparent [5]. However, our work on light field endoscopy reveals several challenges arising from different imaging conditions. In brightfield endoscopic imaging, tissues are opaque; thus, there is always a strong, constant background intensity from illumination of the tissue. Consider that a background signal can be considered as a uniform component $B$ added to a higher-frequency signal $F$ to form an image $I$:

$$I = B + F$$

Under the light field imaging condition, a uniform light source located at infinity would be forward-projected to become a uniform background illumination shared across each elemental image. Thus, placing the background signal at the deepest depths in the reconstructed volume would be a potential solution for the RLD volume. This is confirmed by our experimental results, which show that the intensity increases with depth within the volume, regardless of the target object's true depth. For example, a tilted playing card is placed up to 10.8 mm from the endoscope, yet the volumetric reconstruction shows increased layer intensity at depths far beyond the real position of the playing card surface (**Figure S2**). Towards this end, we added a depth-based regularization to RLD, allowing for some suppression of the accumulation of intensity with increasing depth (**Supplementary Note 1**).

#### Note S4: Microfluidic phantom flow experiment

A microfluidic phantom with square channels ranging in thickness from 0.1 to 1.1 mm was designed, and a positive mold was printed using a micro-3D printer (microArch S140, Boston Micro Fabrication). Then, Sylgard 184 PDMS was prepared at a 1:10 crosslinker:base ratio, vacuum-degassed in a desiccator (Bel-Art) for 2 hours, and poured onto the mold. The system was cured overnight at 60 °C. After curing, the PDMS chip was carefully peeled from the mold. The chip and a standard glass coverslip were placed in a plasma cleaner for 5 minutes to activate the surfaces. Afterward, the glass coverslip was pressed onto the chip with a weight for 10 minutes to seal the microfluidic system. Input and output tubing were inserted into the chip and glued.

A scattering-matched blood phantom was formulated by diluting Intralipid 20% (I141, Sigma Aldrich) into water. Using the reduced scattering coefficient $\mu_s' \cong 1.30\ \text{mm}^{-1}$ of oxygenated blood at 785 nm[15], the desired concentration of Intralipid $v_{il}$ was calculated using a linear relationship [16]:

$$v_{il} = \frac{\mu'_{s,blood}}{\mu'_{s,il}} \qquad (\mathrm{S}-20)$$

Where the reduced scattering coefficient of undiluted intralipid 20% was previously measured to be 19.4 $\text{mm}^{-1}$ at 785 nm [16]. Accordingly, a 6.7% v/v dilution of intralipid 20% was infused into the microfluidic chip as a blood phantom.

During experimentation, a syringe pump (Pump 11 Elite Infuse Only, Harvard Instruments) was used to infuse the microfluidic chip at 5 different volumetric flow rates. A reference RGB image of the microfluidic chip with no flow was first captured through the endoscope. Afterward, the NIR laser illumination was turned on, and speckle video was captured at 10 FPS for 20 minutes with a 10 ms exposure time. During this time, the syringe pump would infuse the chip for 2 minutes, then pause for 2 minutes. Thus, for a single imaging sequence, 5 cycles were completed, with flow rates of 540, 180, 64.8, 27, and 5.4 µL/min for each cycle.

After video capture, raw light field speckle images were processed to produce the flow index and then reconstructed alongside the RGB reference image.

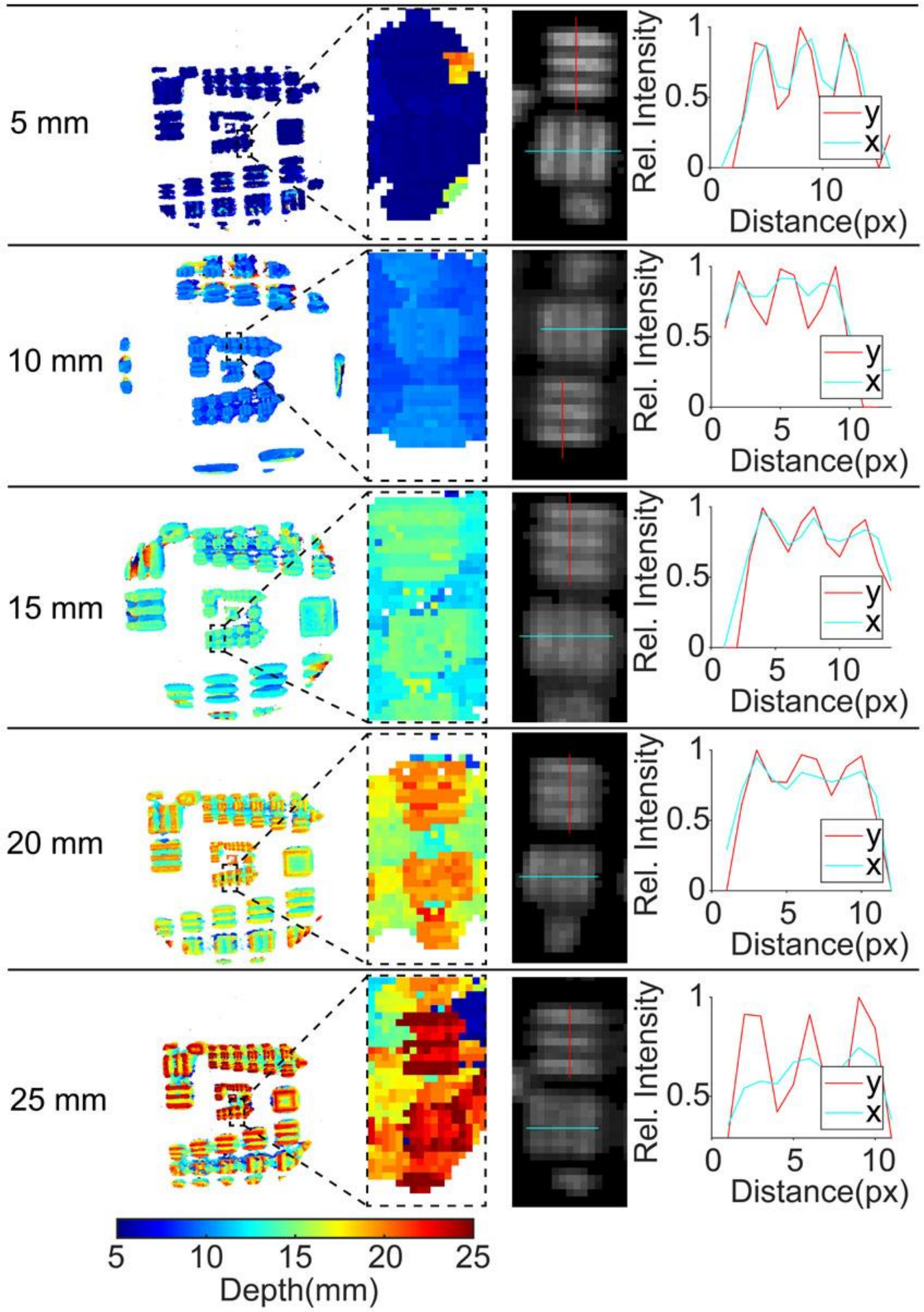


**Figure S1**. USAF target characterization of lateral resolution and depth accuracy for the NIR channel.

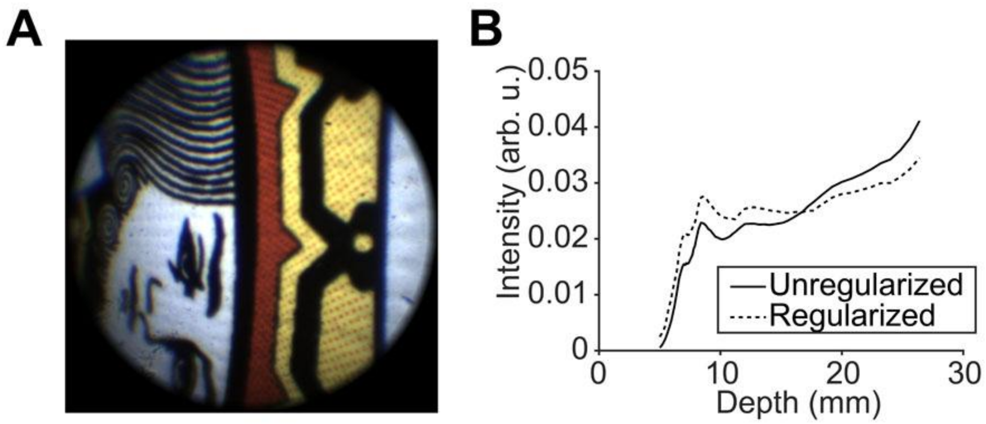


**Figure S2**. Depth vs mean intensity for a tilted playing card spanning depths from 5.6-10.8 mm. (A) Reference RGB central view. (B) Mean intensity profile for volumes reconstructed with and without depth-regularization.

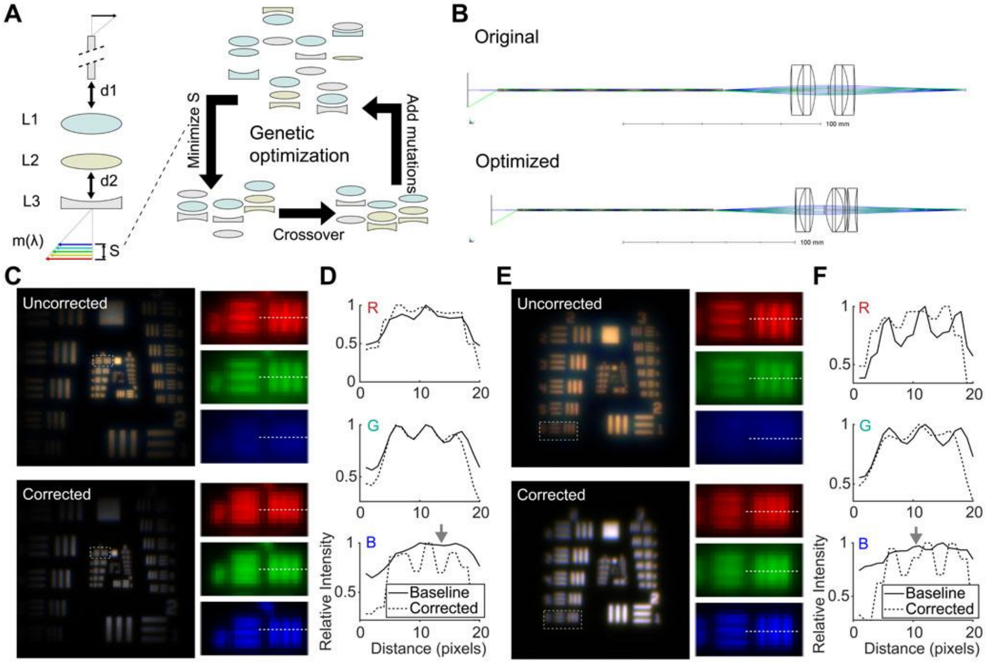


**Figure S3**. Chromatic aberration optimization of the system. (A) Schematic of the system variables that were optimized to yield the minimum range of chromatic focal planes. d, distance. L, lens. m, magnification. S, chromatic focal plane separation (range). (B) Zemax diagrams of the original baseline design compared to the determined optimized design. (C) Experimental comparison of the uncorrected and corrected designs imaging a USAF target at 8 mm distance. (D) Comparison of profiles in (C) for each color channel. (E) Experimental comparison of the uncorrected and corrected designs imaging a USAF target at 18 mm distance. (F) Comparison of profiles for the selected USAF target for each color channel.

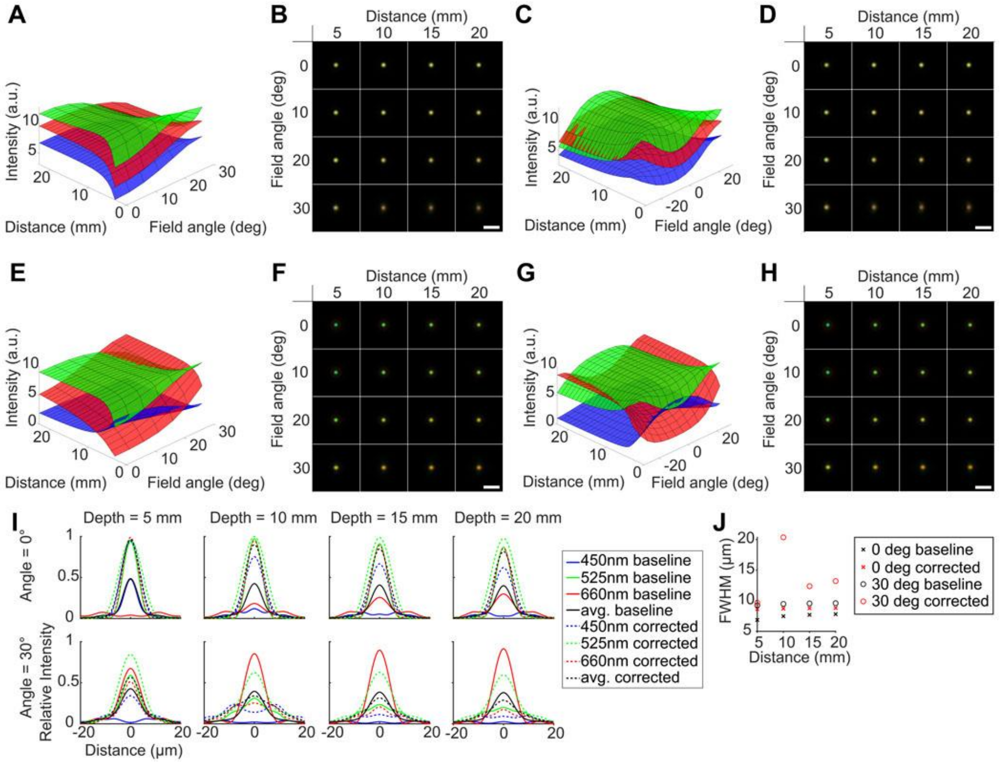


**Figure S4**. Detailed simulation results comparing the PSF of the baseline and optimized system. (A) Optimized design maximum spot intensity for each color channel over field angle and axial depth for the central elemental image. (B) Exemplary RGB PSFs. (C) Baseline design: maximum spot intensity for each color channel over field angle and axial depth in the central elemental image. (D) Exemplary RGB PSFs. (E) Optimized the design maximum spot intensity for each color channel over field angle and axial depth for a peripheral elemental image. (F) Exemplary RGB PSFs. (G) Baseline design max spot intensity for each color channel over field angle and axial depth for a peripheral elemental image. (H) Exemplary RGB PSFs. (I) Comparison of the spot profile for R, G, B, and combined color profiles at different axial depths and 0° to 30° field angle. (J) FWHM of the monochrome color profile for different axial distances at two different field angles. All scale bars: 20 μm.

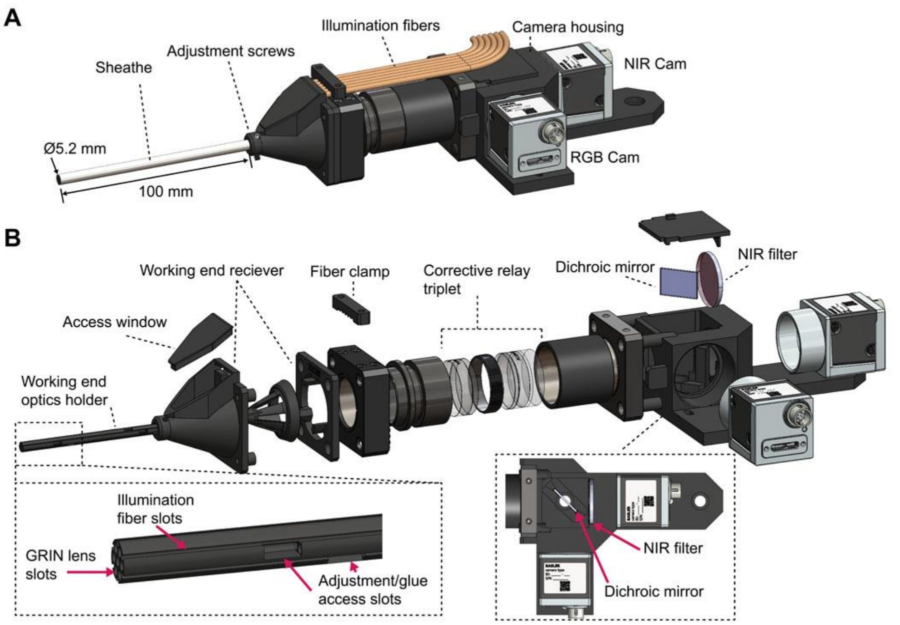


**Figure S5.** Endoscope model. (A) Final design, featuring clinically-relevant working dimensions. (B) Exploded view detailing internal components. Left inset: Close up of the working end optics holder, detailing slots for GRIN optics and fiber illumination. Right inset: top-down view of camera housing, indicating the location of the dichroic mirror and NIR filter to separate image signals.

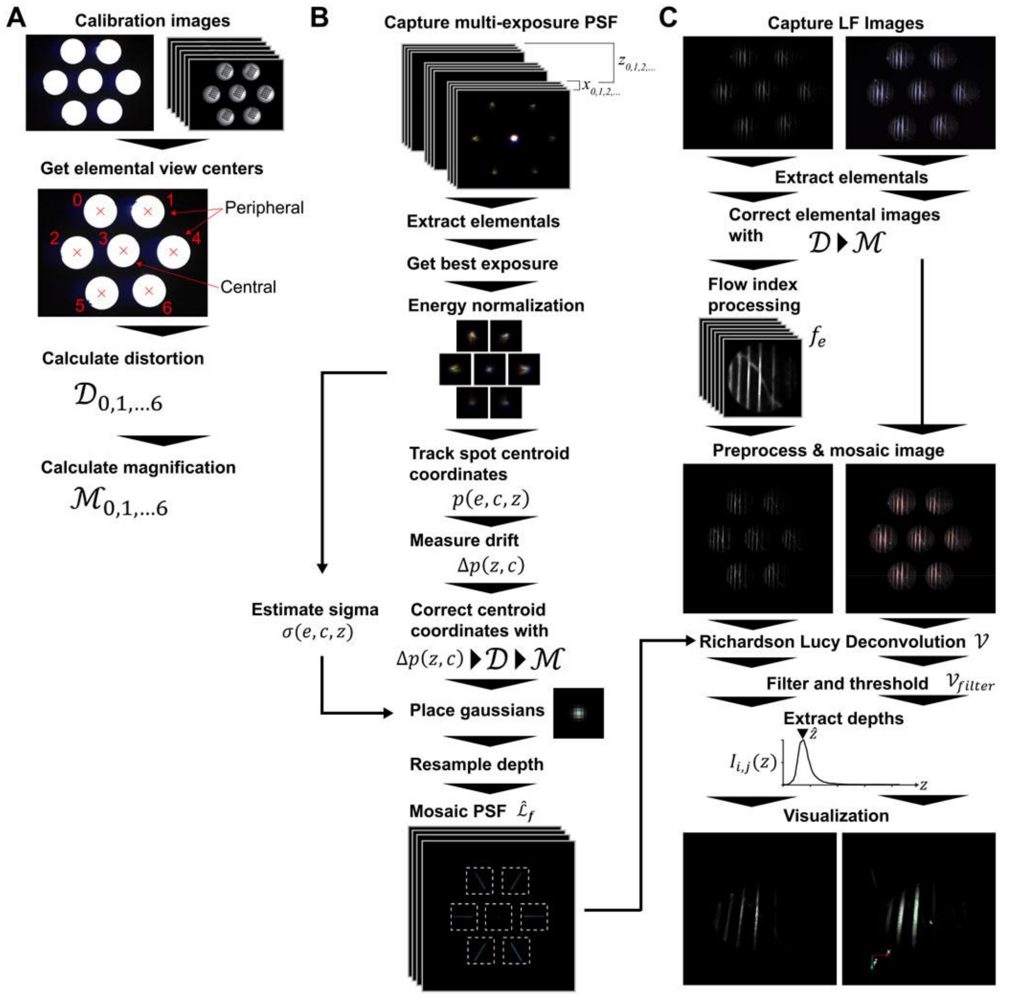


**Figure S6**. Processing pipeline. (A) System correction factors calibration. (B) PSF measurement process. (C) Light field reconstruction processing.

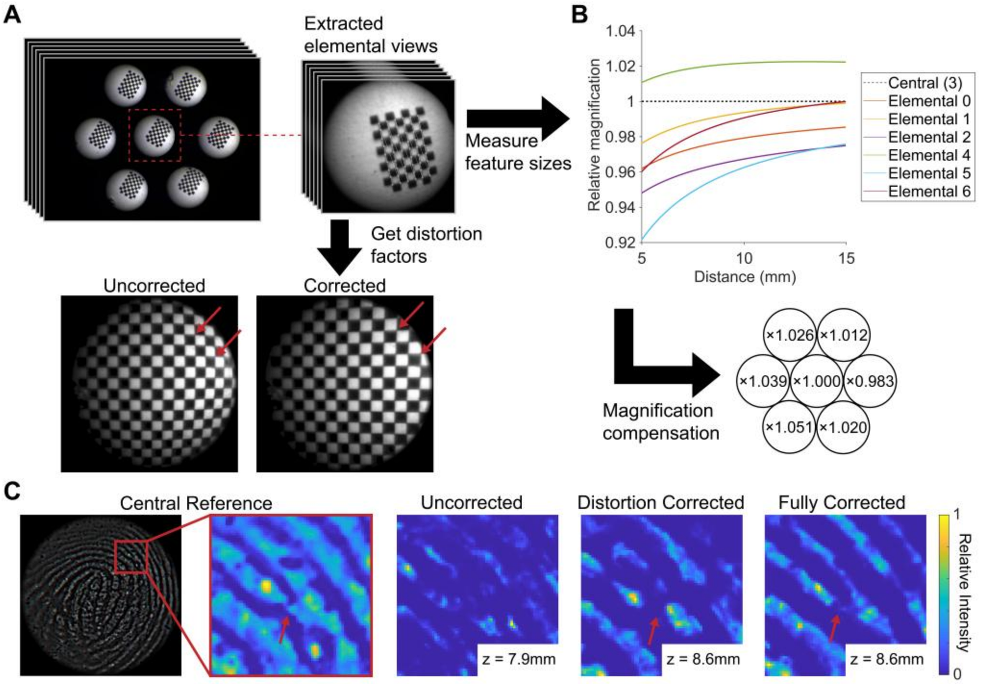


**Figure S7**. Distortion and magnification correction framework. (A) Multiple images of grid features are used to perform distortion correction. The changes in the measured grid-side lengths for each elemental image are also used to compute the relative magnification. (B) Relative magnification of each elemental view with respect to the central elemental image, resulting in a constant magnification compensation factor when averaged. (C) Each calibration step improves the reconstruction quality of the fingerprint. At the focal plane, the bifurcation becomes visible only after both the distortion and magnification correction schemes.

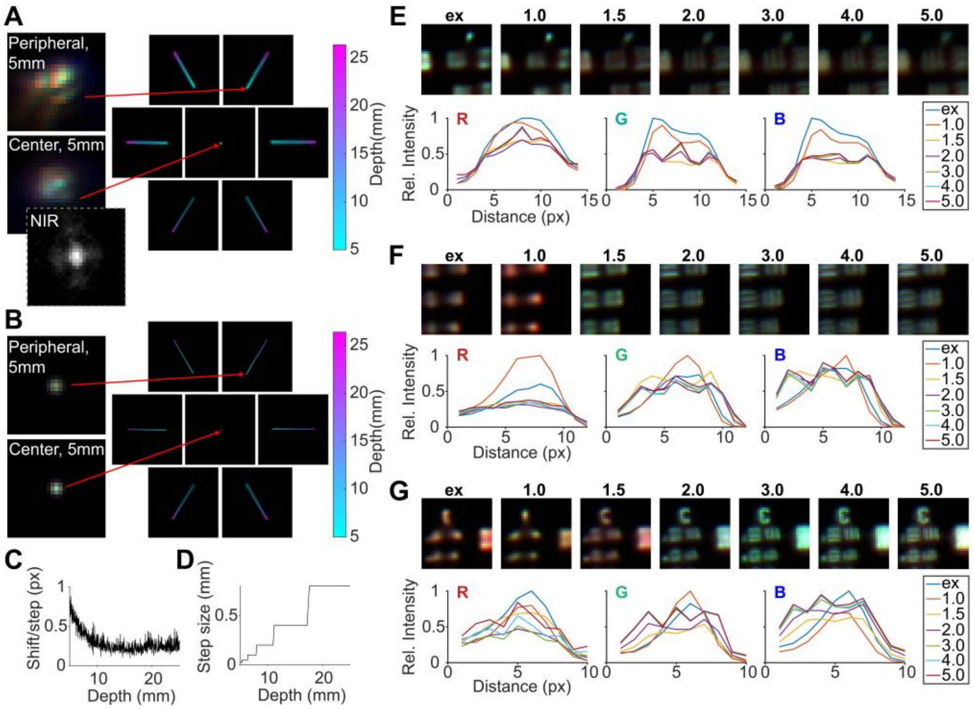


**Figure S8**. PSF visualization, sampling, and hybrid strategy. (A) Left: Close-up experimental RGB PSF spot for a central and peripheral elemental view at 5 mm depth. Inset: Central NIR PSF spot at 5 mm depth. Right: Max projection across the depth of the experimental PSF. Averaged across all color channels (RGB). (B) Left: Close-up contracted hybrid PSF spot for a central and peripheral elemental view at 5 mm depth. Right: Max projection across the depth of the hybrid PSF. Averaged across all color channels (RGB). (C) The magnitude of pixel shift of the centroid of a PSF spot as the depth is incremented by one 25 µm step. (D) Incremental step size over the imaging depth after nonlinear depth resampling. (E) Most in-focus slice of the reconstructed volume of a USAF target at 5 mm deconvolved using the experimental PSF (ex) and different hybrid contraction factors. Graphs: Horizontal intensity profiles of the imaged bar feature for each color channel for each PSF type. (F) Most in-focus slice of the reconstructed volume of a USAF target at 10 mm deconvolved using the experimental PSF (ex) and different hybrid contraction factors. Graphs: Horizontal intensity profiles of the imaged bar feature for each color channel for each PSF type. (G) Most in-focus slice of the reconstructed volume of a USAF target at 20 mm deconvolved using the experimental PSF (ex) and different hybrid contraction factors. Graphs: Horizontal intensity profiles of the imaged bar feature for each color channel for each PSF type.
